# High-throughput functional analysis of CFTR and other apically localized channels in iPSC derived intestinal organoids

**DOI:** 10.1101/2021.07.07.451180

**Authors:** Sunny Xia, Zoltán Bozóky, Onofrio Laselva, Michelle Di Paola, Saumel Ahmadi, Jia Xin Jiang, Amy Pitstick, Chong Jiang, Daniela Rotin, Christopher N. Mayhew, Nicola L. Jones, Christine E. Bear

**Author notes:** Corresponding author at: Hospital for Sick Children, 686 Bay St., Room 20.9420, Toronto, ON, M5G 0A4, Canada.

## Abstract

Induced Pluripotent Stem Cells (iPSCs) can be differentiated into epithelial organoids that recapitulate the relevant context for CFTR and enable testing of therapies targeting Cystic Fibrosis (CF)-causing mutant proteins. However, to date, CF-iPSC-derived organoids have only been used to study pharmacological modulation of mutant CFTR channel activity and not the activity of other disease relevant membrane protein constituents. In the current work, we describe a high-throughput, fluorescence-based assay of CFTR channel activity in iPSC-derived intestinal organoids and describe how this method can be adapted to study other apical membrane proteins. In these proof-of-concept studies, we show how this fluorescence-based assay of apical membrane potential can be employed to study CFTR and ENaC channels and an electrogenic acid transporter in the same iPSC-derived intestinal tissue. This multiparameter phenotypic platform promises to expand CF therapy discovery to include strategies to target multiple determinants of epithelial fluid transport.

## Introduction

There has been remarkable progress made in the use of patient tissue derived primary organoids for the *in-vitro* modeling of Cystic Fibrosis (CF) pathogenesis and testing of therapies targeting mutant CFTR. CFTR mutations (1–5), lead to the loss of CFTR expression and/or function as a phosphorylation-regulated anion channel at the cell surface. Three dimensional (3D) primary organoids have been used effectively to report CFTR-mediated fluid transport as swelling of their luminal cavities (2). Importantly, the rectal organoid model has been shown to recapitulate the genotype specific impact of CF-causing mutations on fluid secretion, while also enabling the ranking of therapeutic interventions targeting defective CFTR expression and function (2, 6). Organoid swelling has been shown to correlate with multiple clinical biomarkers of CF, such as Sweat Chloride Concentration and lung function FEV1 measurements (6). This demonstrates the relevance of patient derived organoids for *in vitro* assessment of patient specific responses to modulators that directly target mutant CFTR.

Luminal swelling is measured in outside-in 3D organoids that enclose the apical membrane in which CFTR is localized. To date, such 3D structures have not been useful for screening the activities of cation channels and electrogenic transporters implicated in net epithelial fluid absorption. In order to solve this problem, 2D monolayer cultures were generated from enzymatically dissociated rectal organoids to provide direct access to the apical membrane (7). While these 2D cultures enabled low-throughput electrophysiological assays of CFTR mediated chloride conductance, they did not reconstitute the native functional expression of ENaC, even though this channel is known to be expressed in the large intestine (7).

Merkert et al, developed 2D intestinal epithelial cultures from CF iPSCs and demonstrated the potential of this model for high throughput drug screening of novel CF therapies (5). In this case, a halide sensitive reporter protein (eYFP) was genetically integrated into *CFTR* to enable studies of CFTR channel activity. However, to date, no similar strategy has been developed for the study of cation channels in iPSC (8).

Given the importance of ENaC and sodium-dependent transporters in modifying net epithelial fluid transport and maintaining epithelial barrier function, there is clearly a need for the development of robust models and assays to test these activities and their modulation. Both of these membrane proteins constitute potential molecular targets for companion therapies to augment the impact of approved CFTR modulators drugs.

In the current work, we describe a method that enables the measurement of CFTR and ENaC activity in *opened* iPSC-derived intestinal organoids, a 2D preparation that retains 3D expression levels of both channels and is adaptable to medium-high throughput, high-content, phenotypic analyses.

## Results

### 3D CF Human Intestinal Organoids (HIO) exhibit defective fluid secretion, which can be restored through the use of CFTR modulators or gene editing

HIOs were differentiated from a homozygous F508del CF-iPSC line, and the isogenic Mutation-Corrected (MC) IPSCs harbouring Wt-CFTR using established protocols (9, 10) (Fig. 1a). Immunostaining confirmed that the HIOs expressed CDX2, E-cadherin, and MUC2, proteins which are characteristic of intestinal epithelial cells (Fig. 1b). Non-CF (MC) iPSCs-derived HIOs exhibited an increase in size after activation by the adenylate cyclase agonist, Forskolin (Fsk), consistent with previously observed CFTR-mediated fluid secretion (2) (Fig. 1c, 1d). CF HIOs generated from iPSCs from a patient homozygous for the major CF-causing mutation, F508del, displayed defective forskolin mediated organoid swelling (Fig. 1c) These results are consistent with the known primary defects conferred by the F508del mutation related to defective CFTR protein processing and function and lack of organoid swelling in primary CF rectal organoids (2). However, the swelling iPSC HIOs organoids exhibits a slower kinetic profile compared to primary rectal organoids (n ≥ 3 biological replicates, n ≥ 50 organoids per biological replicate) (2).

**Figure 1:**
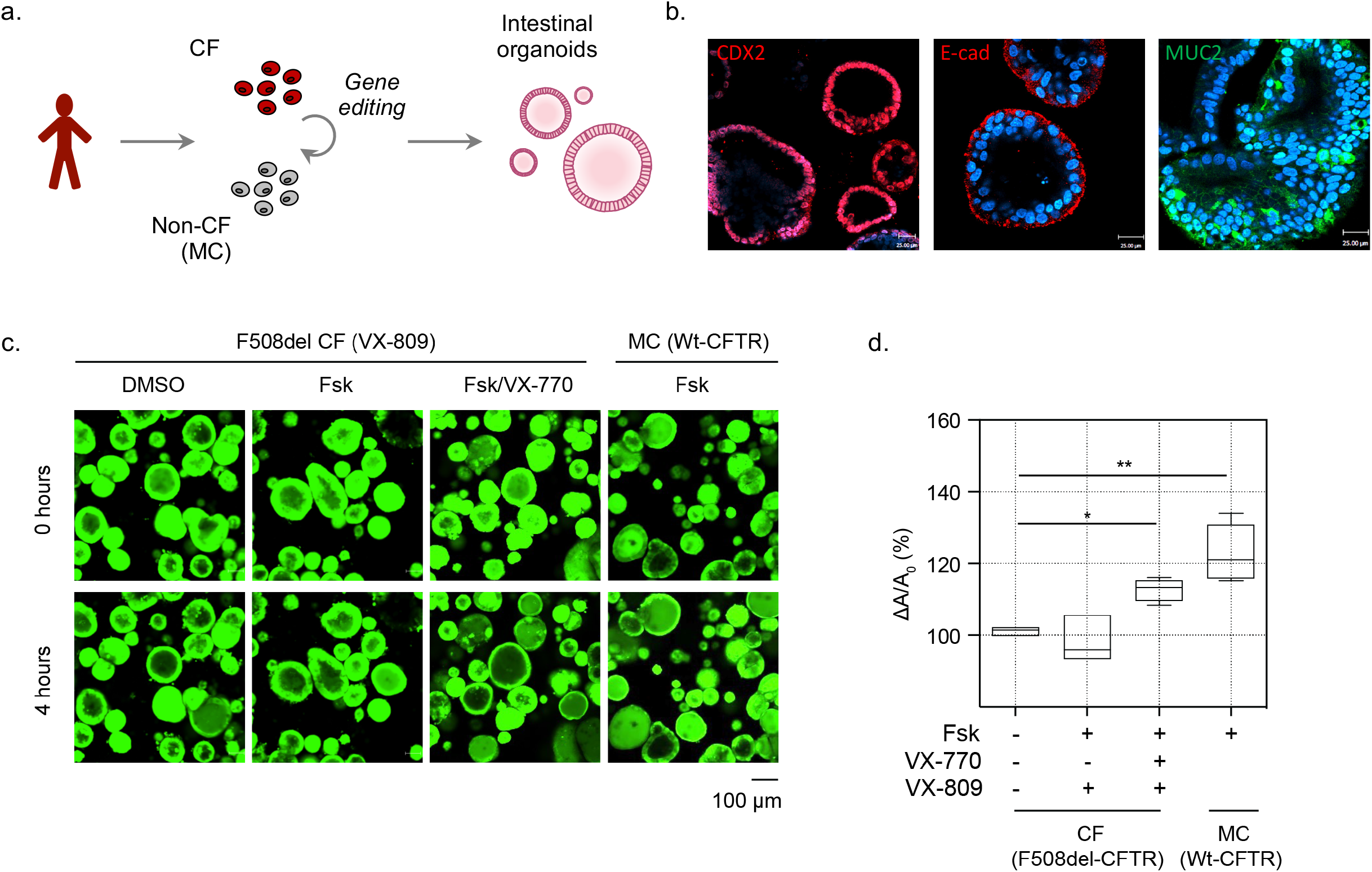
iPSC-derived HIOS can be used to measure function of Wt-CFTR and pharmacologically rescued F508del-CFTR as Fsk induced organoid swelling. **a.** Schematic depicting generation of 3D homozygous F508del CF and isogenic, Mutation Corrected (MC) intestinal organoids from iPSCs. **b.** Characterization of iPSC derived intestinal organoids. Immunofluorescence studies of hPSC derived intestinal organoids highlighting expression of intestinal cell markers CDX2 (red, intestinal marker), E-cadherin (red, epithelial cell), and MUC2 (green, goblet cell). **c.** Representative images of forskolin induced swelling of F508del-CFTR expressing CF organoids or Wt-CFTR expressing Mutations Corrected (MC) organoids. CF organoids were rescued chronically (24 hrs) with DMSO control or VX-809 (3 μM) and acutely stimulated with Fsk (10 μM) or Fsk and VX-770 (1 μM). **d.** Bar graph shows the change in organoid size post Fsk induced swelling (ΔA) relative to average organoid size at baseline (A_0_) (mean ± SEM). (*P = 0.0191, ** P = 0.0065, n ≥ 3 biological replicates. Each biological replicate = independent organoid passage, technical replicate = average of >30 organoids).

In CF organoids, mutant F508del-CFTR protein misprocessing and function (measured as forskolin stimulated swelling) was rescued through pharmacological treatment with CFTR corrector, lumacaftor (VX-809) and acute potentiation with ivacaftor (VX-770) (11). Interestingly, the extent of rescued swelling in F508del CF HIOs was similar to that observed in mutation-corrected, Wt-CFTR expressing HIOs (Fig. 1c, 1d). This result was surprising given the relatively modest rescue effect (approximately 30% of Wt function) induced by this modulator combination in other *in vitro* models (2). Electrophysiological assays of CFTR modulator activity in 2D intestinal monolayers cultures are considered the “gold standard” for testing efficacy. Yet, these methods are relatively low throughput. Thus, we were prompted to develop a complementary assay of CFTR that would enable direct measurement of F508del-CFTR channel modulation in the apical membrane of intestinal organoids, in a high-throughput format.

### *Opened* HIOs enable direct assessment of apical Wt-CFTR channel function in a high-throughput format

As previously demonstrated using primary mouse colonic organoids (12), the removal of the matrigel led to splitting open and access to the apical membrane in 3D organoids. We applied this method to the study of iPSC differentiated HIOs (Fig. 2a). Immunofluorescence studies were conducted to confirm apical membrane location through visualization of tight junction complex protein, Zona Occluden-1 (ZO-1) in the opened iPSC HIOs (Fig. 2b). Further, *opened* organoids can be detected using the FLIPR^®^ membrane potential sensitive fluorescence, which could allow for direct functional assessment of apical membrane protein (Fig. 2c). There were no significant differences in gene expression of CFTR, ENaC, SLC6A14 (an electrogenic amino acid transporter) and other intestinal and epithelial cell type markers in 3D when compared to 2D *opened* organoids (Fig 2d, Supplementary Fig. 1). In addition, we confirmed mature CFTR expression in mutation corrected 3D HIOs and *opened* HIOs (Fig. 2e).

**Figure 2:**
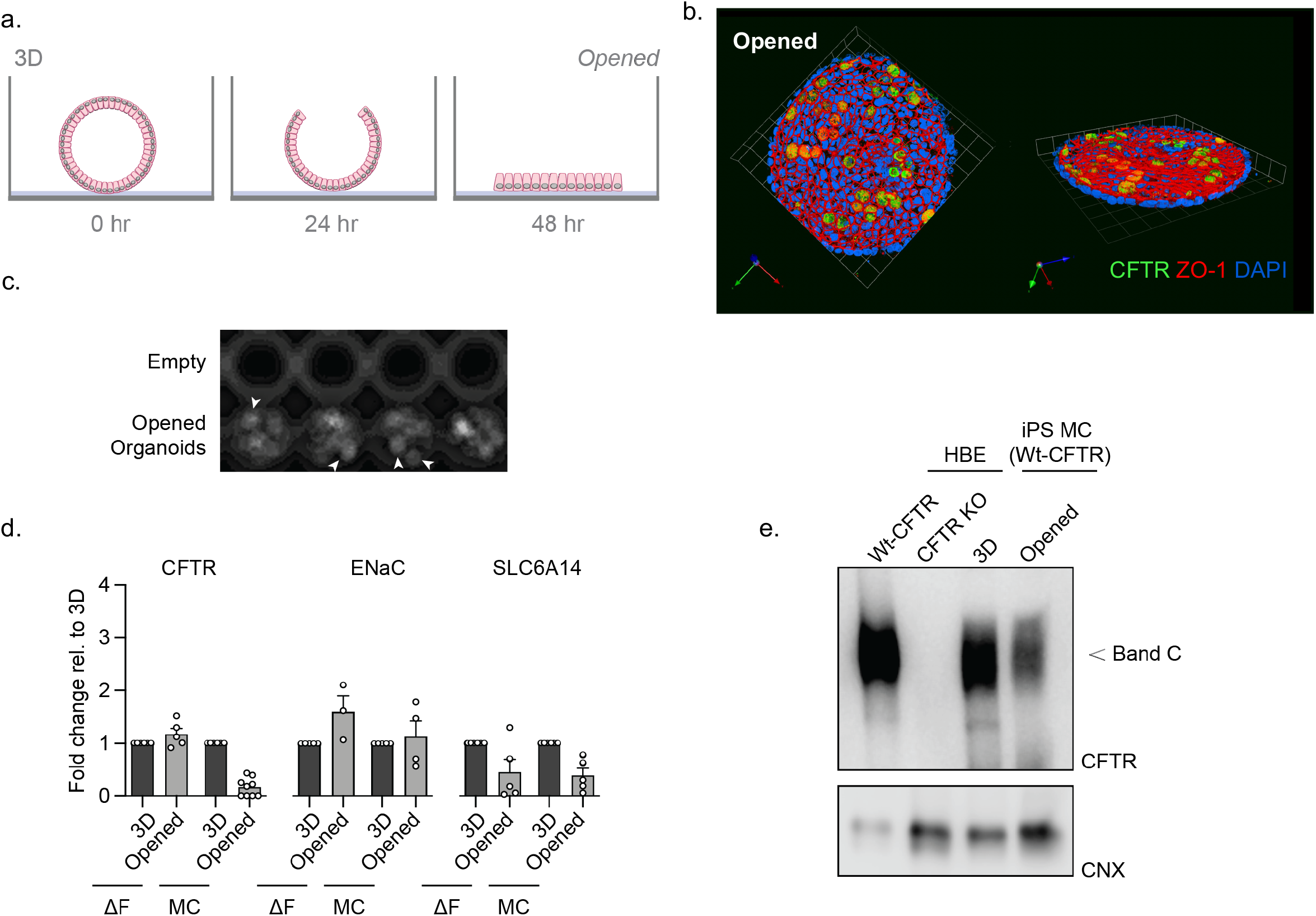
Characterization of *opened* and 3D iPSC-HIOs. **a.** Schematic depicting generation of opened organoids with the removal of the extracellular supporting matrix. **b.** Immunofluorescence of CFTR (green), apical membrane marker, ZO-1(red), and nuclei (blue) in *opened* isogenic non-CF organoids. **c.** Representative raw FLIPR fluorescence image of *opened* organoids, with each object (arrowhead) as an opened organoid. **c.** Gene expression RT-qPCR studies of CFTR and ENaC and SLC6A14 in opened organoids relative to 3D organoid expression. **d.** Western blot of WT-CFTR expression in 3D and opened, mutation corrected organoids, compared to expression in HBE cell line and HBE CFTR knockout cell lines.

After confirmation of CFTR protein expression, CFTR channel function was measured in *opened* HIOs using the Apical Chloride Conductance (ACC) assay (Fig. 3a), as previously show in mouse colonic organoids (12). The MC *opened* differentiated HIOs, which expressed Wt-CFTR, demonstrated remarkably consistent Fsk responses (Fig. 3b). The ACC assay of *opened* MC Wt-HIOs displayed a Fsk dose response with an EC50 of 0.0287 μM (Fig. 3c), which is lower compared to previously reported values in Fsk induced swelling of CF organoids (6). We demonstrated that this assay is scalable to a high-throughput format, supporting its future utility for testing emerging modulators. The *opened* HIOs showed excellent reproducibility of peak response stimulation and consistent activation kinetics with Fsk stimulation and inhibition with CFTRInh-172, reporting a Z’ factor of 0.5294, supporting its utility as a robust assay of dynamic CFTR function in a high throughput format (Fig. 3d, Supplementary Video 1).

**Figure 3:**
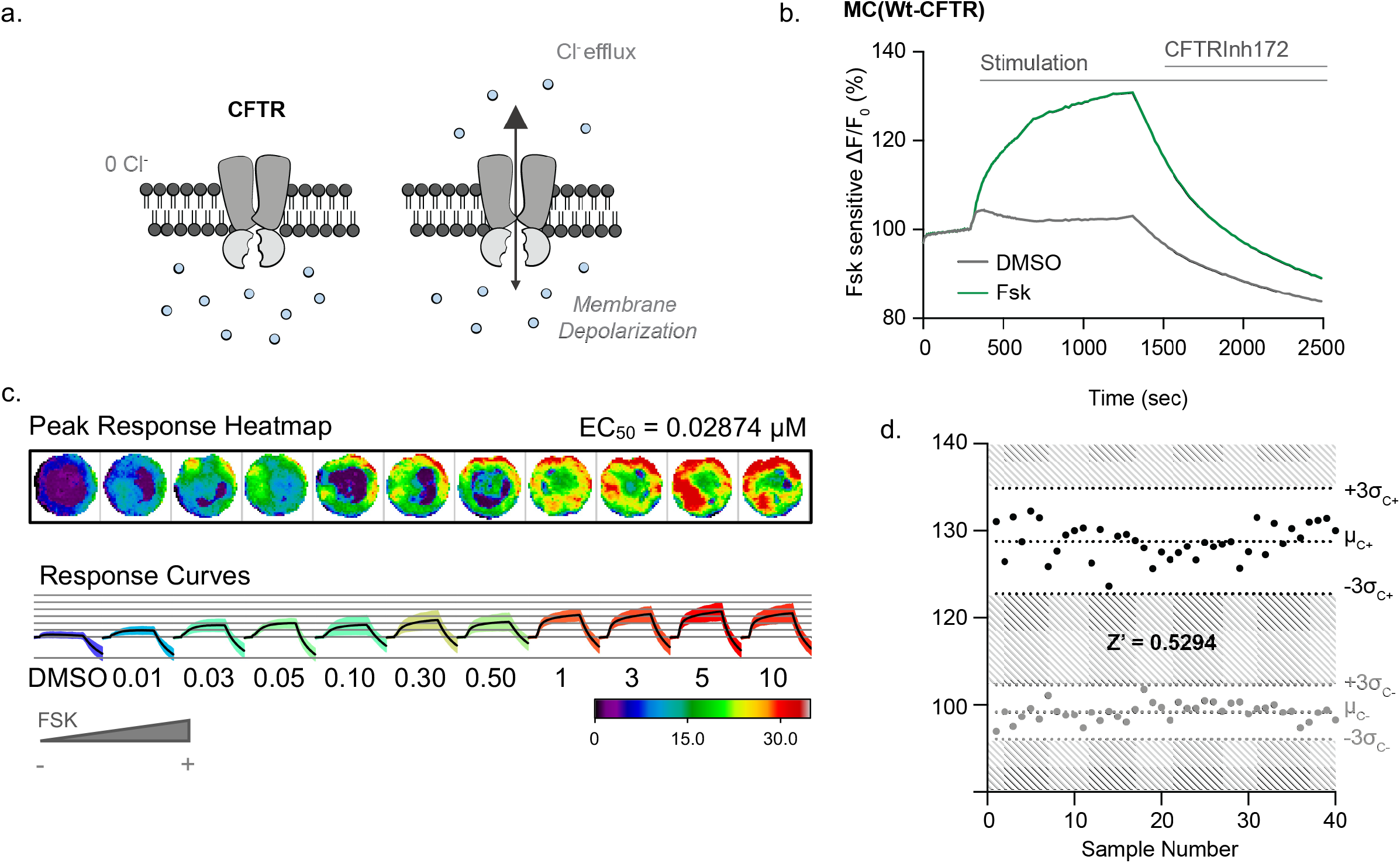
*Opened* iPSC-HIOs enable direct measurements of Wt-CFTR function in a high throughput format. **a.** Schematic depicting the Apical Chloride Conductance (ACC) assay (13). *Opened* MC iPSC-HIOs are placed in a zero-chloride extracellular buffer. Upon addition with Fsk, CFTR mediated chloride efflux leads to increase in membrane potential and the subsequent increase in fluorescence, signal is terminated with acute treatment of CFTRInh172. **b.** Representative trace of Wt-CFTR function measured in MC organoids expressing Wt-CFTR. Opened organoids stimulated with Fsk (10 μM). CFTR response was terminated with CFTRinh172 (10 μM). **c.** Peak responses heatmaps and response curves of opened MC organoids stimulated with increasing concentrations of Fsk. **d.** Bland-Altman plot depicting reproducibility of stimulated CFTR response. Black points measuring maximum change in fluorescence changes with acute Fsk stimulation in comparison to grey points representing DMSO control.

### Opened CF organoids can model pharmacological rescue of F508del-CFTR with CF modulators

We were prompted to determine if the ACC assay is effective in detecting the primary defect caused by the F508del mutation and evaluating the efficacy of clinical modulators on *opened* CF HIOs (Fig. 4a). The *opened* F508del CF HIOs displayed no significant fluorescence changes with Fsk stimulation, consistent with the expected defect in F508del-CFTR channel function prior to modulator rescue (Fig. 4b). VX-809/VX-770 treatment resulted in partial rescue of the mutant F508del-CFTR protein. Furthermore, treatment with the new and highly effective modulator combination, VX-661, VX-445 and VX-770 (TRIKAFTA^TM^) (13), restored F508del-CFTR function to approximately 50% of Wt-CFTR function in MC non-CF HIOs (Fig. 3b-3d).

**Figure 4:**
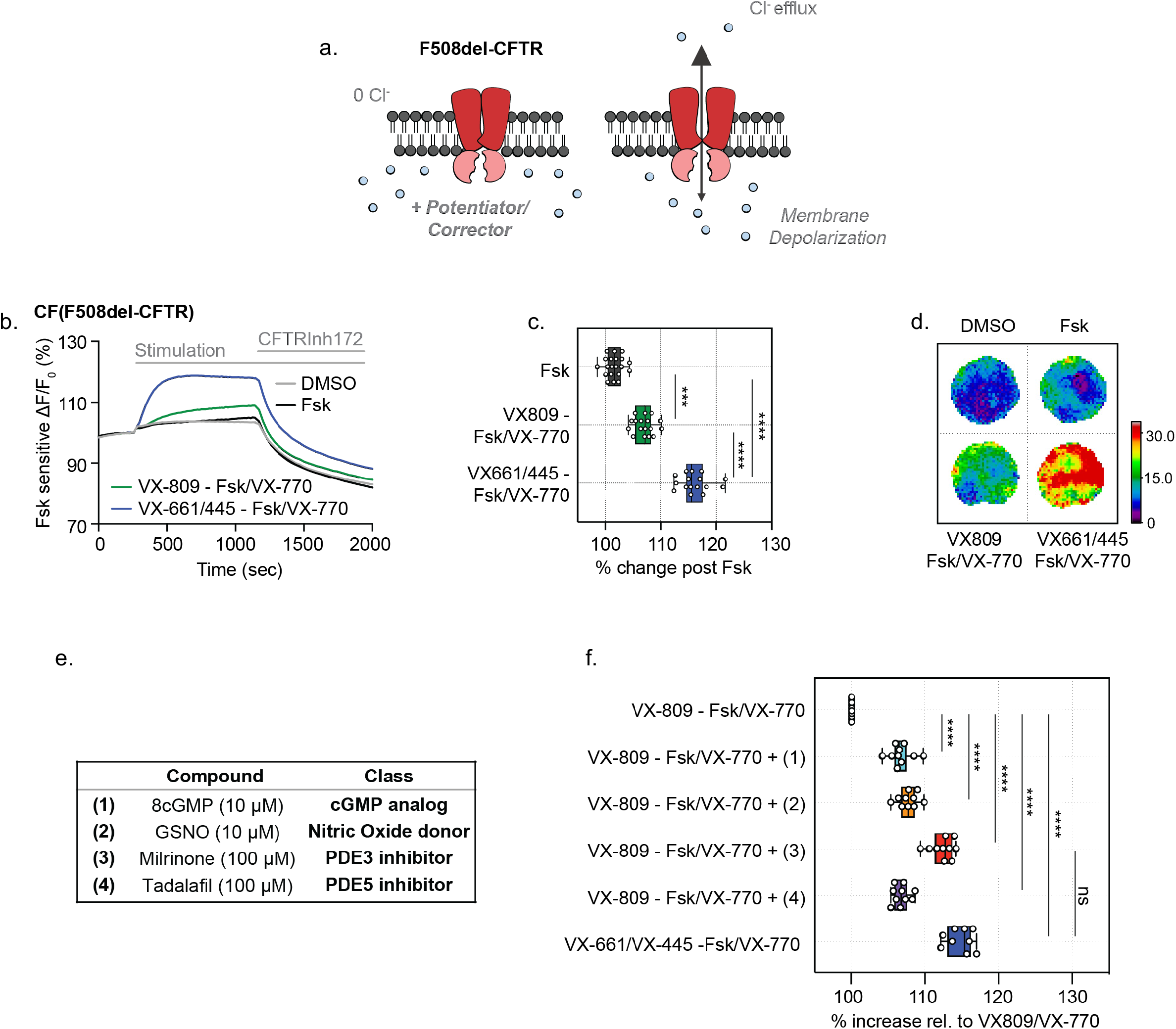
*Opened* CF HIOs can model defective CFTR function and response to CF modulators. **a.** Schematic depicting functional measurement of F508del-CFTR using the ACC assay. **b.** Representative traces of F508del-CFTR response to pharmacological rescue in *opened* CF organoids. *Opened* F508del CF organoids were chronically (24 hr) rescued with VX-809 (3 μM), VX-445 (3 μM)/VX-661 (3 μM) or DMSO as control and acute stimulation with Fsk (10 μM)/VX-770 (1 μM) or Fsk (10 μM)/VX-770 (1 μM). **c.** Box and whisker plot and **d.** Peak response heatmaps of F508del-CFTR response to pharmacological rescue in opened CF organoids. *Opened* iPSC-derived F508del CF organoids were chronically (24 hr) rescued with VX-809 (3 μM), VX-445 (3 μM)/VX-661 (3 μM) or DMSO as control and acute stimulation with Fsk (10 μM) or Fsk (10 μM)/VX-770 (1 μM) (*** P = 0.004, **** P < 0.001, n > 3 biological replicates, n = 3 technical replicates). **e.** Table of compounds tested in combination with Fsk/VX-770 and VX-809. **f.** Box plot shows F508del CFTR stimulation peak response post chronic rescue with VX-809 (3 μM) and acute drug treatment with the listed phosphodiesterase inhibitors and Fsk (10 μM)/VX-770 (1 μM) ****P <0.0001, n = 3 biological replicates, n = 3 technical replicates. Each biological replicate = independent organoid passage, technical replicate = 1 well of 96 well plate).

To determine if iPSC HIOs have the potential to identify companion therapies for CF, we assessed the effects of known modulators of PKA and PKG phosphorylation, since post-translational modifications of CFTR have been implicated in regulating modulator efficacy (Fig. 4e) (14, 15). After partial correction of the trafficking defect in F508del with VX-809, *Opened* F508del CF HIOs were acutely potentiated with Fsk/VX-770 in combination with a nitric oxide (NO) agonist targeting enhancement of the PKG phosphorylation pathway (8cGMP or GSNO), which have been shown to augment VX-809 rescued F508del-CFTR activity (16, 17). Alternatively, *opened* F508del CF HIOs were treated with phosphodiesterase inhibitors (Milrinone or Tadalafil), which have been shown to be effective in stimulation of F508del-CFTR short circuit current in murine intestinal tissue (18). Milrinone addition, along with VX-809/VX-770, significantly increased the Fsk response to levels comparable to the triple modulator combination (VX-661/VX-445/VX-770) treatment (Fig. 4e-4f). Therefore, the ACC assay is sufficiently sensitive to distinguish between various modulator combinations that are expected to exhibit different efficacies in rescuing the functional expression of F508del-CFTR. With direct access to the apical membrane of HIOs in the *opened* format, this prompted us to determine whether functional output of other apical membrane channels can be detected. Since ENaC is functionally expressed in the intestinal epithelium (19), we tested the utility of the measuring ENaC mediated changes membrane potential in a high-throughput format.

### Measurement of ENaC specific activity in MDCK overexpression cells

We first developed an assay measuring ENaC function using an engineered cell line in which the three ENaC subunits were stably expressed. The renal epithelial MDCK cells, which is genetically engineered to express HA tagged-αENaC, myc (& T7) tagged-βENaC, flag tagged-γENaC and the un-transfected parental MDCK cell line (20) (Fig 5a). In order to measure constitutive ENaC function, we established an inward sodium gradient and assessed the effect of the ENaC inhibitors, amiloride and phenamil on the apical membrane of confluent differentiated, MDCK monolayers. Under these conditions, we predict that the apical membrane potential as monitored by the novel Apical Sodium Conductance (ASC) assay, would hyperpolarize upon inhibition of ENaC with amiloride or phenamil (Fig. 5b). As expected, MDCK cells expressing tagged αβγENaC, exhibited membrane hyperpolarization after addition of either amiloride or phenamil at 10 or 50 μM concentrations in a sodium dependent manner (Fig. 5c, 5d, Supplementary Fig. 2). These responses were significantly greater in the MDCK cells expressing tagged αβγ-ENaC than in the parental line.

**Figure 5:**
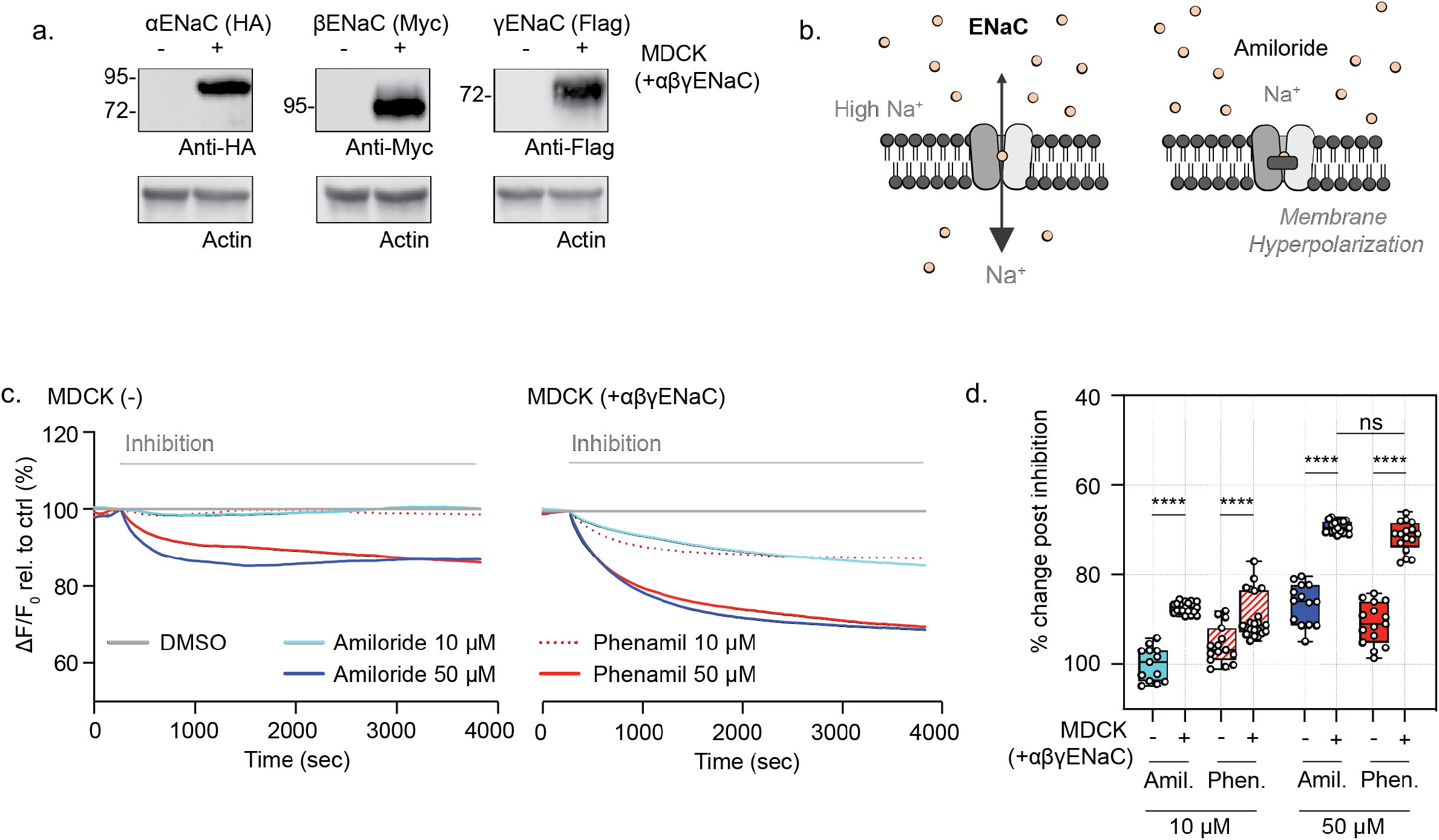
Validation of ENaC function in ENaC over expression MDCK cell line. **a.** ENaC subunit expression in MDCK cells that are stably transfected compared to parental (untransfected) MDCK control cells. Western blot detection of αENaC, βENaC, or γENaC, with anti-HA, anti-Myc or anti-flag antibodies, respectively. **b.** Schematic depicting ENaC inhibition in the novel Apical Sodium Conductance assay (ASC). In presence of the high extracellular sodium, acute addition of amiloride and amiloride analogues result in ENaC inhibition and relative membrane hyperpolarization, which is detected as decrease fluorescence signal. **c.** Representative traces and **d.** box and whisker plot of ENaC inhibition in MDCK cells and MDCK cells over expressing triple epitope tagged αβγENaC with amiloride and amiloride analogue, phenamil amiloride (10 μM and 50 μM) relative to DMSO control (**** P > 0.0001, n = 4 biological replicates, n ≥ 4 technical replicates).

### Function of ENaC and the sodium dependent transporter, SLC6A14 can be measured*opened* CF and Mutation Corrected (MC) HIOs

We were then prompted to determine if ENaC function was measurable in opened HIOs using the assay developed above. Similar to the response measured in MDCK cells expressing ENaC, we found that amiloride (10 uM) addition evoked a hyperpolarization response in opened HIOS (Fig. 6a, 6c). Furthermore, this response in *opened* HIOS was recapitulated using the amiloride analogue, phenamil (Fig. 6c) (21). Similar to ACC assay of CFTR channel function (Fig. 2d), the ASC based assay of ENaC function in *opened* iPSC differentiated HIOs showed excellent reproducibility and consistent amiloride response leading to a Z’ factor of 0.573. Such parameters indicate the iPSC HIO model and ASC assay together provide an excellent candidate platform for monitoring dynamic and high throughput drug screening of potential ENaC modulators (Fig. 6b, Supplementary Video 2), further validating the suitability of the ASC *Opened* HIOs for drug screening and evaluation of modulator efficacy (Fig. 6c).

**Figure 6:**
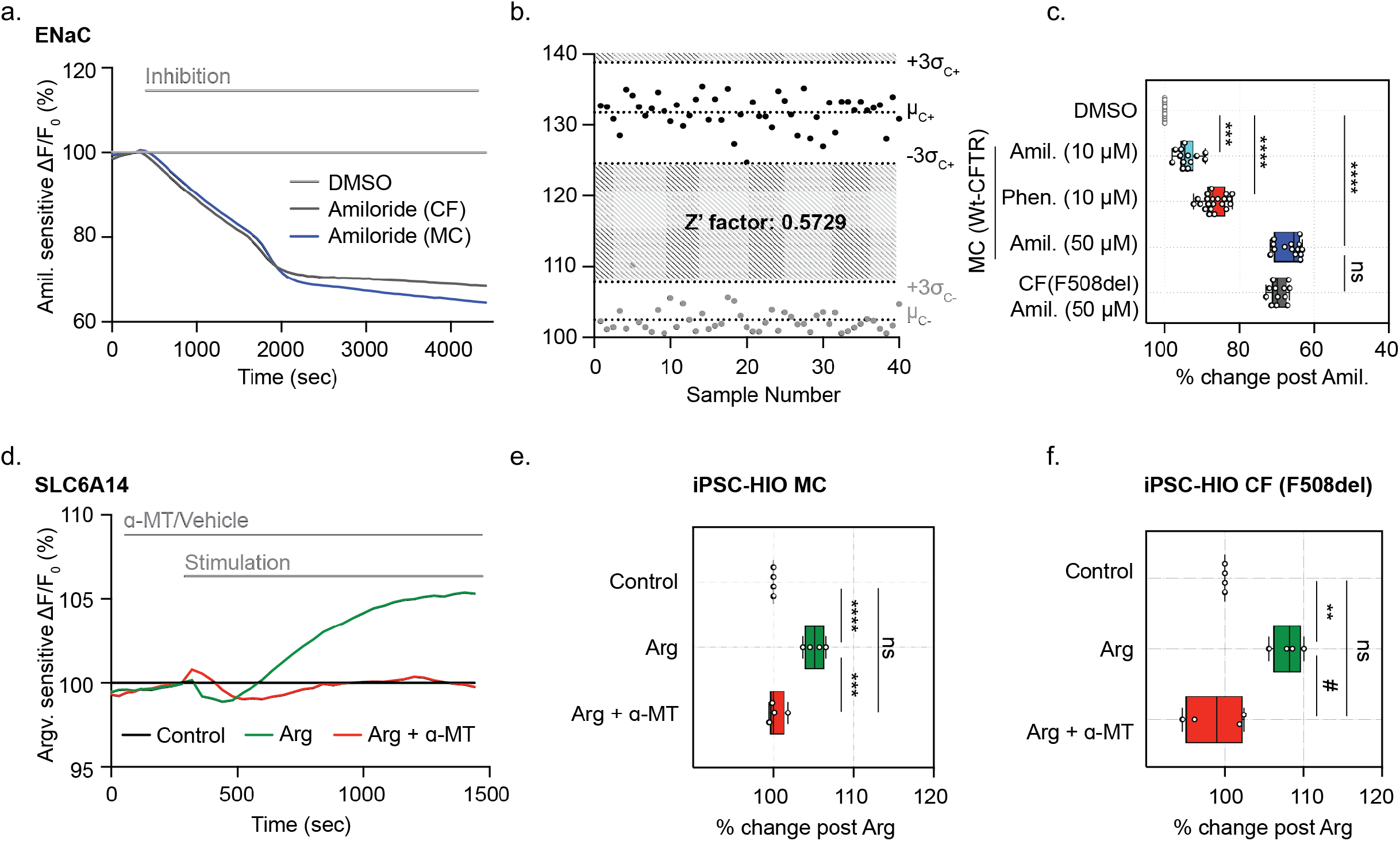
Opened iPSC differentiated HIOs enables high throughput assessment of ENaC specific function, response to ENaC modulators and measurement of SLC6A14 activity. **a.** Representative ENaC inhibition in *opened* F508del CF and MC organoids with acute treatment with Amiloride (50 μM). **b.** Bland-Altman Plot depicting the reproducibility of ENaC inhibition with amiloride treatment. Black points measuring maximum inhibition in fluorescence changes with acute amiloride treatment in comparison to grey points representing DMSO control. **c.** Box and whisker plot shows ENaC inhibition with acute treatment with amiloride (10 μM and 50 μM), or Phenamil, (all 10 μM) in opened iPSC MC HIOs organoids (*** P = 0.0002, **** P > 0.0001, n > 3 biological replicates, n > 3 technical replicates). **d.** Representative trace of SLC6A14 activity in opened iPSC MC HIOs organoids pretreated with either vehicle control or specific inhibitor, α-MT (2 mM). Opened CF and MC organoids were acutely treated with Arg (1 mM). **e.** Box and whisker plot of SLC6A14 function with acute treatment with Arg (1 mM) in iPSC MC HIOs. SLC6A14 activity was inhibited with pretreatment of specific inhibition α-MT (2 mM) (*** P = 0.0001, **** P > 0.0001, n = 4 biological replicates, n = 3 technical replicates). **f.** Box and whisker plot shows SLC6A14 activity with acute treatment with Arg (1 mM) or in presence of specific inhibitor α-MT (2 mM) in iPSC CF HIOs (** P = 0.0080, # P = 0.0023, n = 3 biological replicates, n = 3 technical replicates).

Previously, the *opened* organoid model was applied to murine intestinal organoids in order to determine the activity of the sodium dependent amino acid transporter, SLC6A14 (12). SLC6A14 is a sodium and chloride dependent electrogenic amino acid transporter expressed in the airway, intestinal, and colonic epithelial tissues. SLC6A14 mediates the uptake of cationic and neutral amino acids along with two sodium ions and one chloride ion, generating one net positive charge translocation and membrane depolarization per amino acid transport (12, 22). Through application of a low sodium/chloride extracellular gradient, SLC6A14 mediated depolarization could also be measured in *Opened* iPSC differentiated HIOs (Fig. 6d). In the presence of low extracellular sodium and chloride, the addition of arginine (L-Arg) to the apical surface of these *opened* organoids evokes apical membrane depolarization in both CF and MC HIOs. As expected on the basis of previous studies, this signal was abolished by the addition of the SLC6A14 blocker, α-Methyl-DL-tryptophan (α-MT) (Fig. 6d-6f) (22). The stimulation with acute L-Arg treatment and specific absence of stimulation with L-Arg and α-MT suggests SLC6A14 amino acid uptake function can be measured using *opened* HIOs.

## Discussion

Stem cell-derived organoids have been employed to advance CF therapy development (2, 4, 6). Evidence for a positive effect of CFTR modulatory compounds on luminal swelling by CF patient-derived primary rectal organoids has been proposed as a potential diagnostic tool to inform personalized medical treatment. In the current work, we described novel methods for studying ion channels and transporters in the apical membrane of patient-specific, iPSC-derived intestinal organoids. In contrast to the widely used, organoid swelling assay of CFTR channel function and previously described 2D monolayer models (2, 7), opened organoids enable the functional measurement of electrogenic apical membrane proteins beyond CFTR using a fluorescence-based assay of membrane potential. We described how the membrane potential sensitive dye, FLiPR can be used to measure function of apical CFTR and ENaC channels as well as the electrogenic amino acid transporter, SLC6A14. Hence, multiple, disease-relevant, electrogenic membrane proteins can be interrogated in the same patient-derived tissue. With these innovations, we have expanded the potential application of CF organoid models to include therapy testing for multiple therapeutic targets.

In addition to its potential application to the study of multiple apical membrane constituents, this assay system is suitable for high-throughput screening and potential drug discovery. The Apical Chloride Conductance (ACC) and Apical Sodium Conductance (ASC) assays of patient-derived tissues exhibit excellent reproducibility with Z’ factor scores of 0.529 and 0.573, respectively. Because of the capacity of this platform for profiling multiple small molecule combinations simultaneously, we found that phosphodiesterase inhibitors, could be used as a companion therapy in combination with ORKAMBI™ to boost F508del-CFTR chloride channel activity to levels comparable to that achieved by the new triple modulator combination, TRIKAFTA™. Therefore, the *opened* organoid can serve as a tool in identifying alternative therapeutics for patients with limited access to TRIKAFTA™. Likewise, the *Opened* organoids also enabled the first evaluation of ENaC modulators in a high-throughput manner in human intestinal tissue. Since the functional expression of multiple channels and transporters can be detected in the *opened* organoid model, this provides the potential to investigate the coordinated regulation of SLC6A14, CFTR, and ENaC, studies that are not feasible in the closed 3D system (23). Together, the *opened* organoids have the potential for identifying potential ENaC modulators in a high-throughput format which can be further investigated and characterized using electrophysiological ussing studies.

Our previous studies have demonstrated SLC6A14 amino acid transporter function is able to augment Wt-CFTR and F508del-CFTR channel function in primary mouse colonic organoids (12). Here, we show that SLC6A14 is functionally expressed in both iPSC differentiated CF F508del and MC *opened* organoids. SLC6A14 has been shown in mouse colonic organoids to mediate F508del-CFTR fluid secretion function through uptake of L-Arg leading to activation of the Nitric Oxide pathway (12). Hence, future studies can focus on studying the potential impact of modulators of SLC6A14 on the functional rescue of F508del-CFTR to mediate CF intestinal disease.

iPSCs have the potential for the differentiation of multiple CF-affected tissues, including the airways, intestines, bile duct and pancreas (4, 9, 24–26). In our proof-of-concept studies, the opened organoids enabled the functional output measurement of multiple membrane channels in iPSC differentiated *opened* CF HIOs and mutation corrected, Wt-CFTR expressing, isogenic HIOs. Patient derived iPSCs provides the opportunities for simultaneous *in vitro* multi-tissue differentiation from individual CF patients to interrogate tissue specific disease pathologies.

In summary, we demonstrated the ability to detect the function of multiple apical membrane proteins in human tissues in a format suitable for in-depth analysis of ion channel regulation and interaction. The high-content and high-throughput capacity of this format will facilitate progress in understanding the impact of the membrane protein context on normal and mutant CFTR channel function.

## Methods and Materials

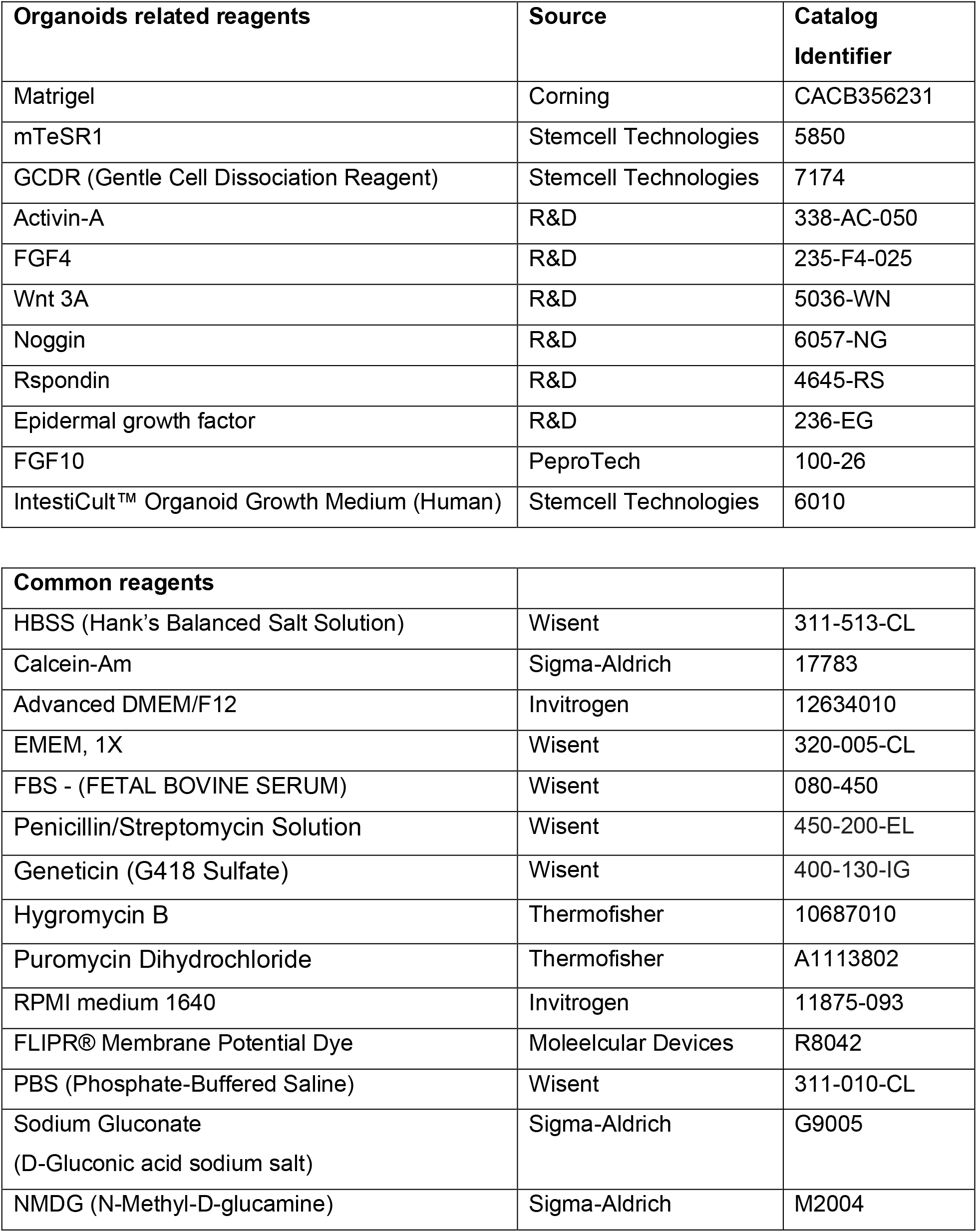
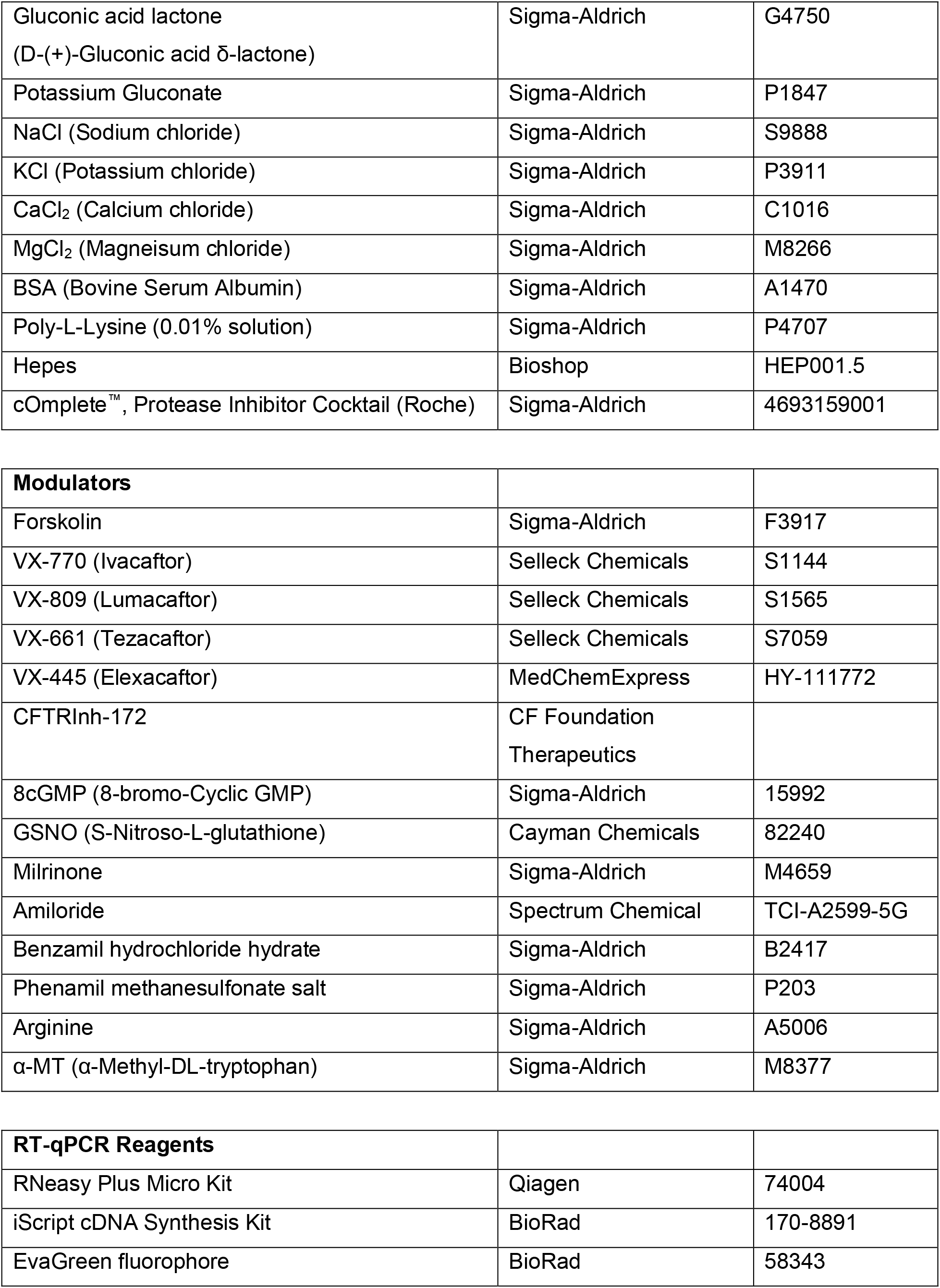
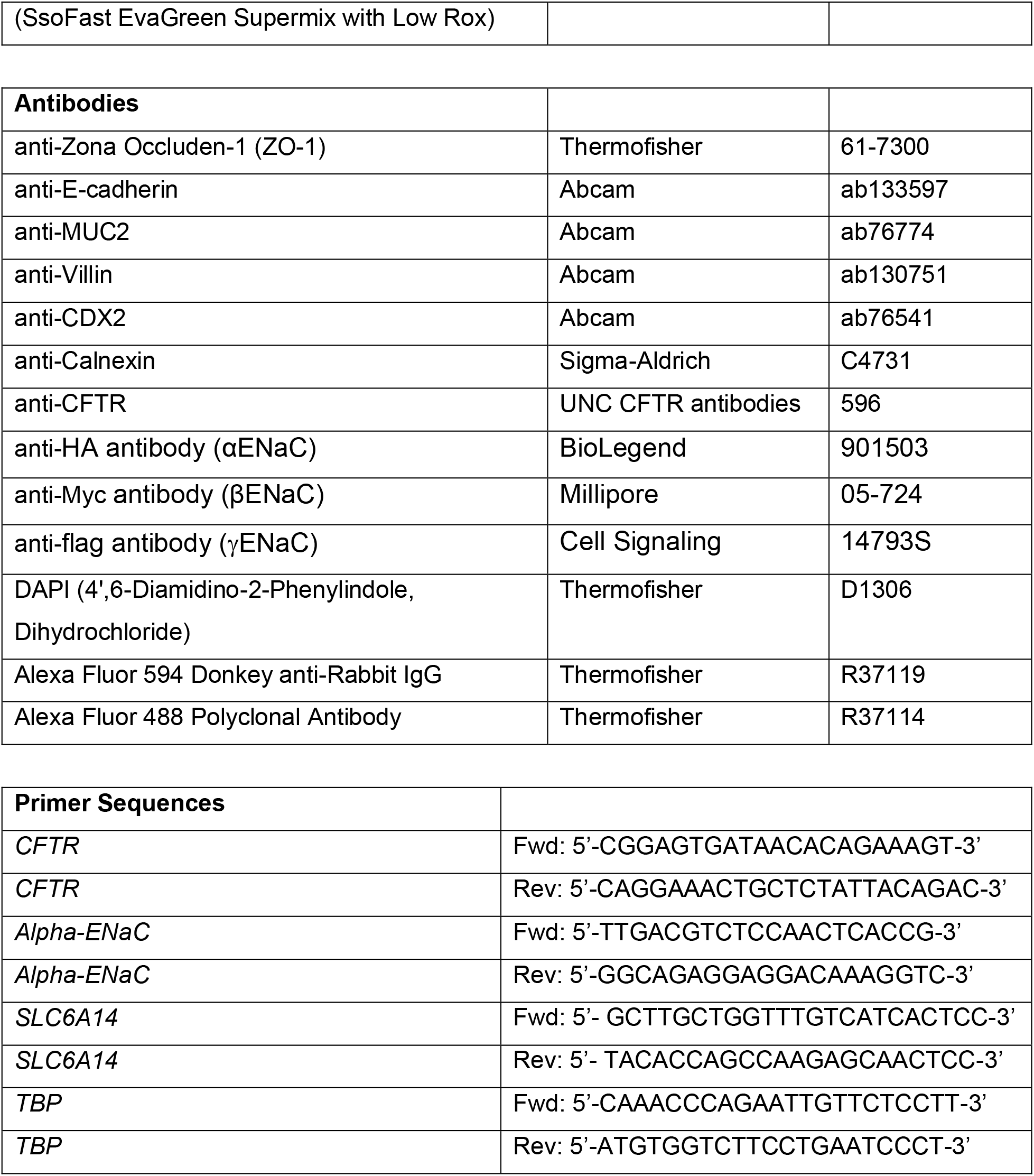

### Cell culture

#### iPSC intestinal organoid

Human intestinal organoids were differentiated as previously described (10). In brief, iPSCs were cultured on ESC qualified-Matrigel^®^ coated 24 well plates in mTeSR1 media. At approximately 60% confluency, differentiation to definitive endoderm was initiated through addition of Activin-A (100 ug/uL), for 3 days. Cultures were then exposed to hindgut endoderm differentiation media containing FGF4 (500 ng/mL) and Chiron99021 (3 uM) for 4 days. Post hindgut differentiation, budding immature organoids were collected and embedded into 50 uL solid Matrigel^®^ drops. Spheroids were cultures for 30 days in previously established growth factor conditioned media (2). Organoids were passaged every 7-10 days and media was changed once every 3 days. To passage organoids, organoids were first collected in ice cold PBS and pelleted through centrifugation for 5 min at 300*g*, 4°C. Post centrifugation, excess PBS, Matrigel^®^ and cellular debris was aspirated. The pelleted organoids were re-suspended in 1 mL of GCDR and incubated at room temperature for 5 mins. With a P1000 pipettor, the organoids were fragmented through pipetting 40-60 times. Organoid fragments were then pelleted through centrifugation and re-suspended in fresh Matrigel^®^ domes and seeded at a 1:3 ratio. Growth factor conditioned medium was added to after allowing the Matrigel^®^ to solidify at 37°C for 35 mins.

#### MDCK and MDCK (tagged αβγENaC) cells

As previously described (20), both MDCK and MDCK (HA tagged-αENaC, myc tagged-βENaC, flag tagged-γENaC) cells are grown in DMEM media, supplemented with 10% FBS, 1% Penicillin/Streptomycin Solution and additional selection antibiotics (300μg/ml G418, 100μg/ml Hygromycin B, 2μg/ml Puromycin). MDCK cells are cultured in presence of Amiloride (10 μM). 24 hours prior to ASC assay, αENaC expression in MDCK cells was induced with dexamethasone (1 μM) and sodium butyrate (10μM) to enhance protein expression of all ENaC subunits.

### Swelling assay

iPSC derived organoids were isolated from the Matrigel^®^ support using ice cold HBSS (Hank’s Balanced Salt Solution) and pelleted through centrifugation. The cell pellet was re-suspended in HBSS containing 3 μM of live cell maker dye, Calcein-AM at 37°C for 45 mins. Excess dye was removed through centrifugation and organoids were resuspended in fresh HBSS for mouse colonic organoids or DMEM for human intestinal organoids. Organoid swelling was induced using Fsk at different concentrations as mentioned or in combination with CFTR potentiators. Mouse colonic organoid swelling was tracked for 30 mins and imaged at 5 mins intervals using fluorescence microscopy (Nikon Epifluorescence/Histology Microscope). Human intestinal organoid swelling as tracked for 4 hours and imaged at 15 mins intervals using confocal microscopy (Nikon A1R Confocal Laser Microscope). Organoid swelling analysis was performed using Cell Profiler v3.1.

Swelling images were analyzed using an in house developed algorithm. In brief. Images were exported as individual TIFF files and aligned using translation registration by cross-correlation. A histogram derived thresholding method (triangle) was used to identify specific organoids in the images. The center of the object masks was used to track individual organoids along the experiment. The differential organoid size at each time point was calculated by subtracting the size of the initial timepoint. Organoids that were not picked over the entirety of the swelling time course were excluded.

### *Opened* organoid cultures

Organoids were removed from the Matrigel^®^ domes and collected in ice cold Advanced DMEM and pelleted through centrifugation. Pellets were then resuspended in growth factor conditioned medium and plated onto Poly-L-Lysine (0.01% solution) coated 96 well plates. Plates were coated following manufacture instructions. Media was changed one day post seeding. All functional studies were done two days after organoid plating.

### Membrane potential based functional assays

#### Apical Chloride Conductance (ACC) Assay for CFTR function

The ACC assay was used to assess CFTR mediated changed in membrane depolarization using methods as previously described (16). In summary, split open primary rectal and iPSC derived intestinal organoids were incubated with zero sodium, chloride and bicarbonate buffer (NMDG 150 mM, Gluconic acid lactone 150 mM, Potassium Gluconate 3 mM, Hepes 10 mM, pH 7.42, 300 mOsm) containing 0.5 mg/ml of FLIPR^®^ dye for 30 mins at 37°C. Wt-CFTR function in iPS cell derived gene edited organoids and primary rectal non-CF organoids was measured after acute addition of Fsk (10 μM) or 0.01% DMSO control. In iPS cell derived F508del CF organoids and primary rectal F508del organoids, cells were chronically rescued with corrector compounds for 24 hours (VX-809 (3 μM), VX-445/VX-661 (both, 3 μM), or DMSO control). Post drug rescue, F508del-CFTR function was measured after acute addition of Fsk (10 μM) and VX-770 (1 μM). Additional modulators (8cGMP, GSNO, Milrinone, and Tadalafil) were added during the FLIPR^®^ dye loading process. CFTR functional recordings were measured using the FLIPR^®^ Tetra High-throughput Cellular Screening System (Molecular Devices), which allowed for simultaneous image acquisition of the entire 96 well plate. Images were first collected to establish baseline readings over 5 mins at 30 second intervals. Modulators were then added to stimulate CFTR mediated anion efflux. Post drug addition, CFTR mediated fluorescence changes were monitored and images were collected at 15 second intervals for 70 frames. CFTR channel activity was terminated with addition of Inh172 (10 uM) and fluorescence changes were monitored at 30 second intervals for 25 frames.

#### Apical Sodium Conductance (ASC) Assay for ENaC function

The ASC assay was used to assess ENaC inhibition upon amiloride addition through assessing changes in membrane hyperpolarization. Split open primary rectal and iPS derived intestinal organoids were incubated with a physiological sodium gluconate buffer (Sodium Gluconate 150mM, Potassium Gluconate 3mM, Hepes 10 mM, pH 7.42, 300 mOsm), containing FLIPR^®^ dye 0.5 mg/ml for 30 mins at 37°C (27). After dye loading, the plate was transferred to the FLIPR^®^ Tetra High-throughput Cellular Screening System (Molecular Devices). Baseline readings were acquired for 5 mins at 30 sec intervals. In iPS cell derived MC organoids, ENaC inhibition was measured following acute addition of Amiloride (50 μM or 10 μM), Benzamil (10 μM), Phenamil (10 μM) or DMSO control. In primary rectal CF and non-CF organoids, ENaC inhibition was measured with acute addition of Amiloride (50 μM) or DMSO control. ENaC mediated membrane hyperpolarization was tracked over time as loss in fluorescence signal over 70 mins at 60 sec intervals.

#### Apical Amino Acid Conductance (AAC) Assay for SLC6A14 function

The AAC assay was used to measure SLC6A14 mediated acute uptake of Arginine leading to membrane depolarization. iPS cell derived gene edited organoids and F508del CF organoids (rescued chronically for 24 hours with VX-809, or DMSO control) were incubated in with a low sodium, low chloride buffer (NMDG 112.5 mM, Gluconic acid lactone 112.5 mM, NaCl 36.25 mM, Potassium Gluconate 2.25 mM, KCl 0.75 mM, CaCl_2_ 0.75 mM, MgCl_2_ 0.5 mM, and HEPES 10 mM, pH 7.42, 300 mOsm), containing FLIPR^®^ dye 0.5 mg/ml for 40 mins at 37°C (17). During dye loading, organoids were treated with α-MT (2 mM), or buffer control. After dye loading, the plate was transferred to the FLIPR^®^ Tetra High-throughput Cellular Screening System (Molecular Devices) (22). Baseline readings were acquired for 5 mins at 30 sec intervals. Arginine (1 mM) was added acutely and change in fluorescence was recorded at 30 sec intervals. SLC6A14 function in primary non-CF rectal organoids was measure with acute addition of Arginine (1 mM) and fluorescence measurements were collected as described above.

#### Analysis and heatmap generation

Experiments were exported as multi frame TIFF images of which every frame recorded the entire plate. Pixels outside of well areas were filtered out using the initial signal intensities and wells containing opened organoids were separated. All traces were normalized to the last point of the baseline intensity. Peak response for each pixel was calculated as the maximum deviation from baseline. During the stimulation segment, fluorescence intensity increased for CFTR and SLC6A14 function, and decreased for ENaC function. Heatmap representation was generated from the peak response of each pixel and the mean response trace of wells was generated by averaging the corresponding pixel traces.

### Real-time Quantitative PCR

As previously described (28), organoid samples were collected in ice cold Phosphate Buffered Saline and pelleted through centrifugation. Total mRNA from pelleted samples was extracted using RNeasy^®^ Plus Micro Kit, following enclosed instructions. After measuring the spectrophotometric quality of extracted RNA through 260/280 ratios of 2.0and 260/230 ratios of 1.8-2.2, mRNA samples with concentrations greater than 300 ng/μL were used to reverse transcribe 1 μg of cDNA using iScript™ cDNA Synthesis Kit. Expression levels of target genes were measured using primers listed above using the FX96 Touch™ Real-Time PCR Detection System using the SYBR Green Master Mix containing the EvaGreen^®^ fluorophore.

### Immunofluorescence

Samples were fixed and permeated with 100% methanol at −20° C for 10 mins. Post methanol incubation, samples were washed 3 times with PBS, 5 mins per wash at room temperature. Following the washes, samples were blocked using 4% BSA for 30 mins and incubated with primary antibody against Zona Occluden-1 (ZO-1) overnight. After removal of primary antibody, samples were wash 3 time with PBS, 5 mins per wash and incubated with secondary antibodies and nuclear marker DAPI for 1 hour. Samples were then washed 3 times with PBS, 5 mins per wash at room temperature. Confocal imaging was done using Nikon A1R Confocal Laser Microscope.

### Western blotting

Samples were collected in ice cold PBS and pelleted through centrifugation at 4°C (500*g* for 7 mins). Post centrifugation, the cell pellet was re-suspended in 200μL of modified radioimmunoprecipitation assay butter (50 mM Tris-HCl, 150 mM NaCl, 1 mM EDTA, pH 7.4, 0.2% (v/v) SDS and 0.1% (v/v) Triton X-100) containing a protease inhibitor cocktail for 10 min. After centrifugation at 13,000 rpm for 5 min, the soluble fractions were analyzed by SDS-PAGE on 6% Tris-Glycine gel. After electrophoresis, proteins were transferred to nitrocellulose membranes and incubated in 5% milk and CFTR bands were detected using the mAb 596. Calnexin (CNX) was used as a loading control and detected using a Calnexin-specific rAb (1:5000). The blots were developed with using the Li-Cor Odyssey Fc (LI-COR Biosciences, Lincoln, NE, USA) in a linear rage of exposure (1-20 min). Relative levels of CFTR protein were quantitated by densitometry of immunoblots using ImageStudioLite (LI-COR Biosciences, Lincoln, NE, USA).

## Supporting information

videos for phenotypic assays

## Acknowledgements

We would like to thank Jacqueline McCormack and Amy P Wong for their help in editing the manuscript. We would also like to thank Christopher Fladd and SPARC BioCentre at the Hospital for Sick Children for help in conducting the organoid swelling assays.

## Author Contributions

S.X., S.A., and C.E.B. conceptualization and experimental design. S.X., B.Z., O.L., M.D., and C.J. performed experiments and data analysis. J.J., and A.P. cultured and differentiated iPCS organoids. S.X., and C.E.B. wrote the manuscript. R.D., C.N.M., N.L.J., and C.E.B. reviewed and revised the manuscript. All authors have read and agreed to the published version of the manuscript.

## SUPPLEMENTARTY

**Supplementary Figure 1:**
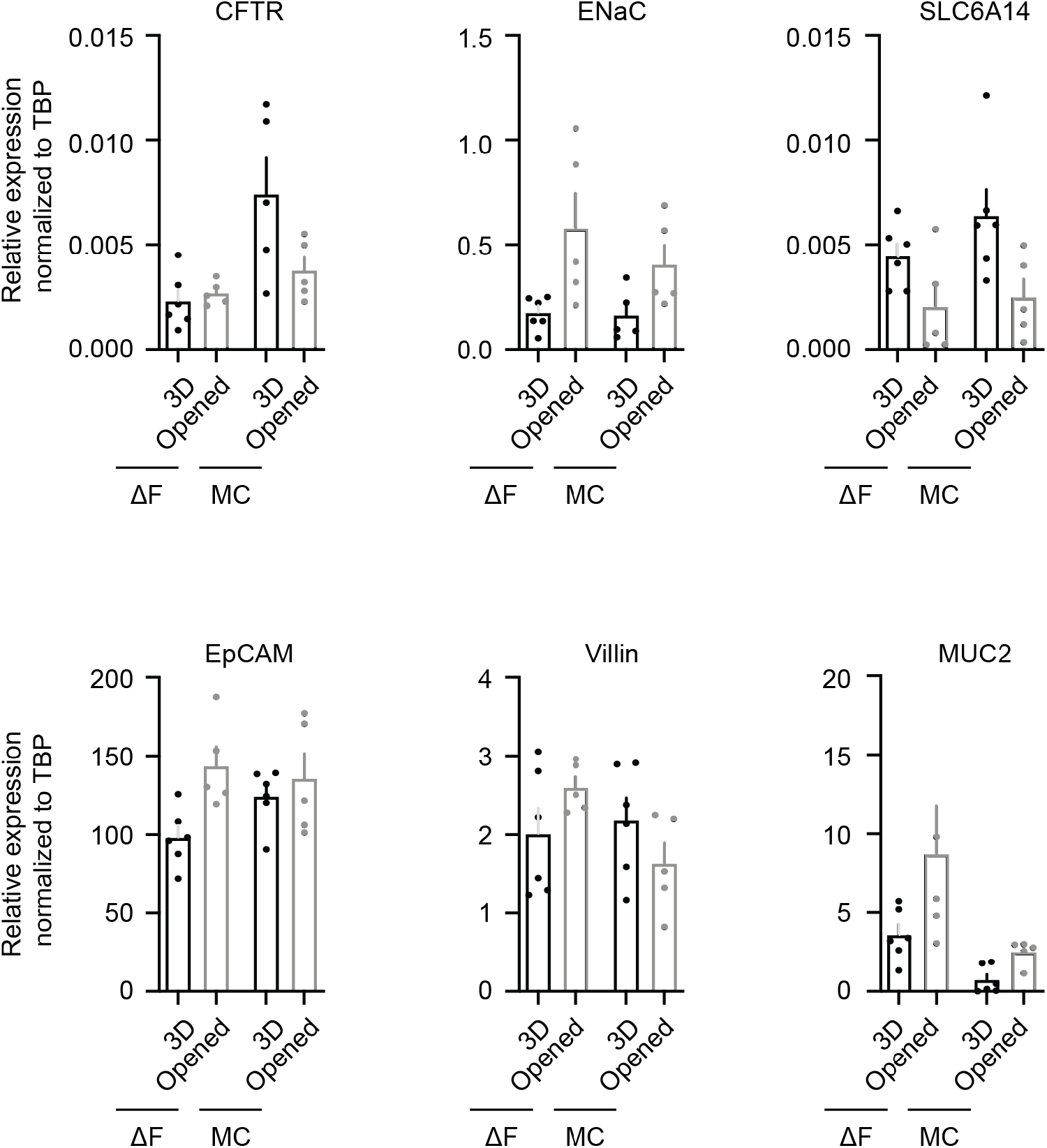
Gene expression studies on iPS derived HIOs. Expression of intestinal apical membrane ion channels (CFTR and ENaC), amino acid transporter (SLC6A14), epithelial cell marker (EpCAM), intestinal epithelial cell marker (Villin) and goblet cells marker (MUC2), relative to house keeping gene *TBP*, in CF and MC organoids in 3D and opened formats using RT-qPCR.

**Supplementary Figure 2:**
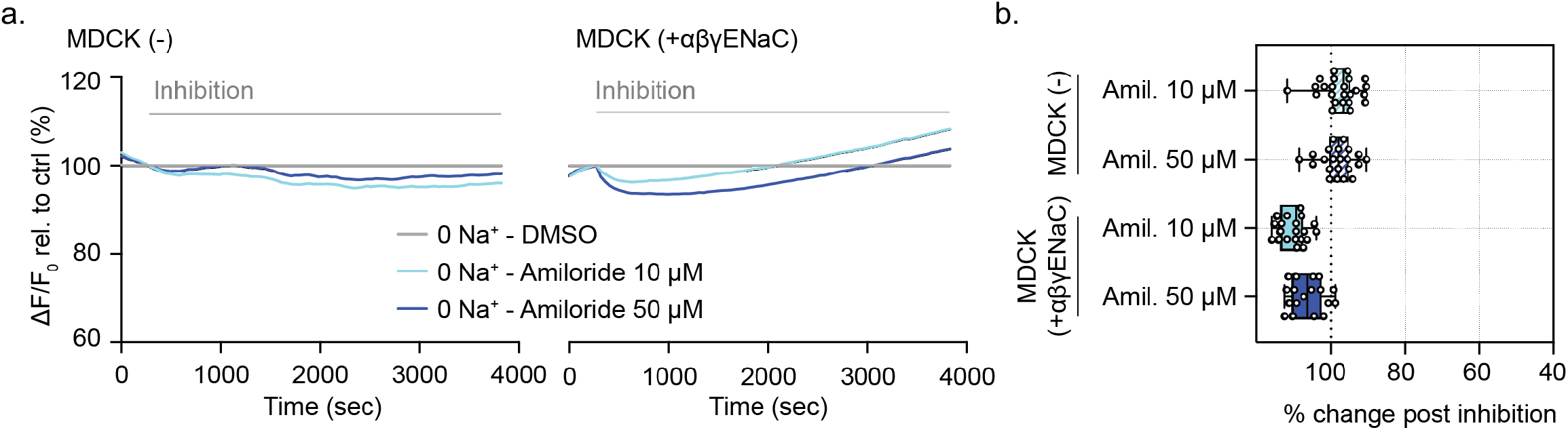
Functional validation of ENaC activity measured in MDCK. a. Representative traces and b. box and whisker plot of parental MDCK control cells or MDCK cells expressing αβγENaC acutely treated with amiloride (10μM or 50μM) in presence and absence of 140mM extracellular sodium (**** P < 0.001, n > 3 biological replicates, n = 3 technical replicates).

