## Supplementary material for "High-throughput functional analysis of CFTR and other apically localized channels in iPSC derived intestinal organoids": videos for phenotypic assays

### Slide 1
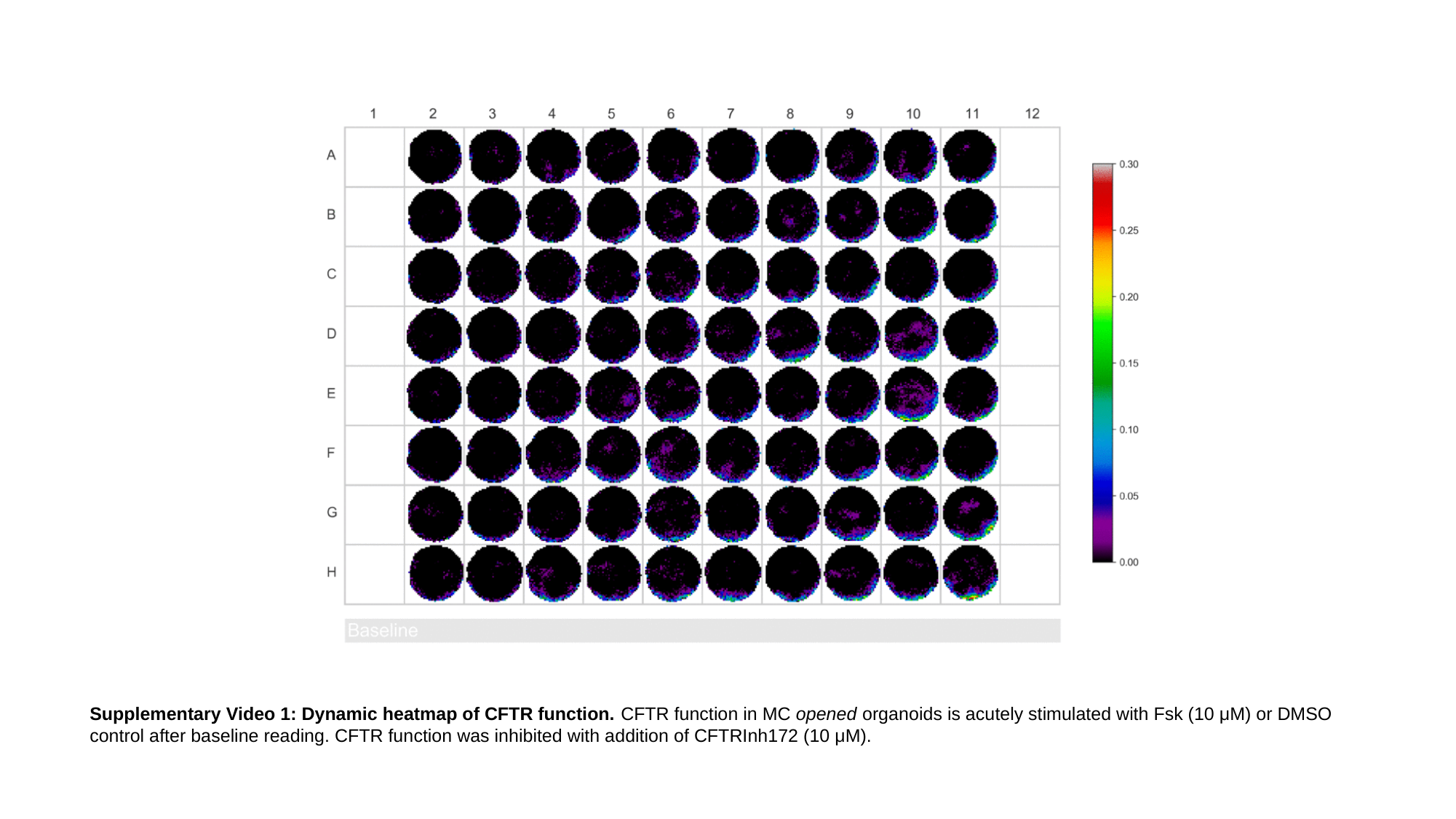

Supplementary Video 1: Dynamic heatmap of CFTR function. CFTR function in MC opened organoids is acutely stimulated with Fsk (10 μM) or DMSO control after baseline reading. CFTR function was inhibited with addition of CFTRInh172 (10 μM).

### Slide 2
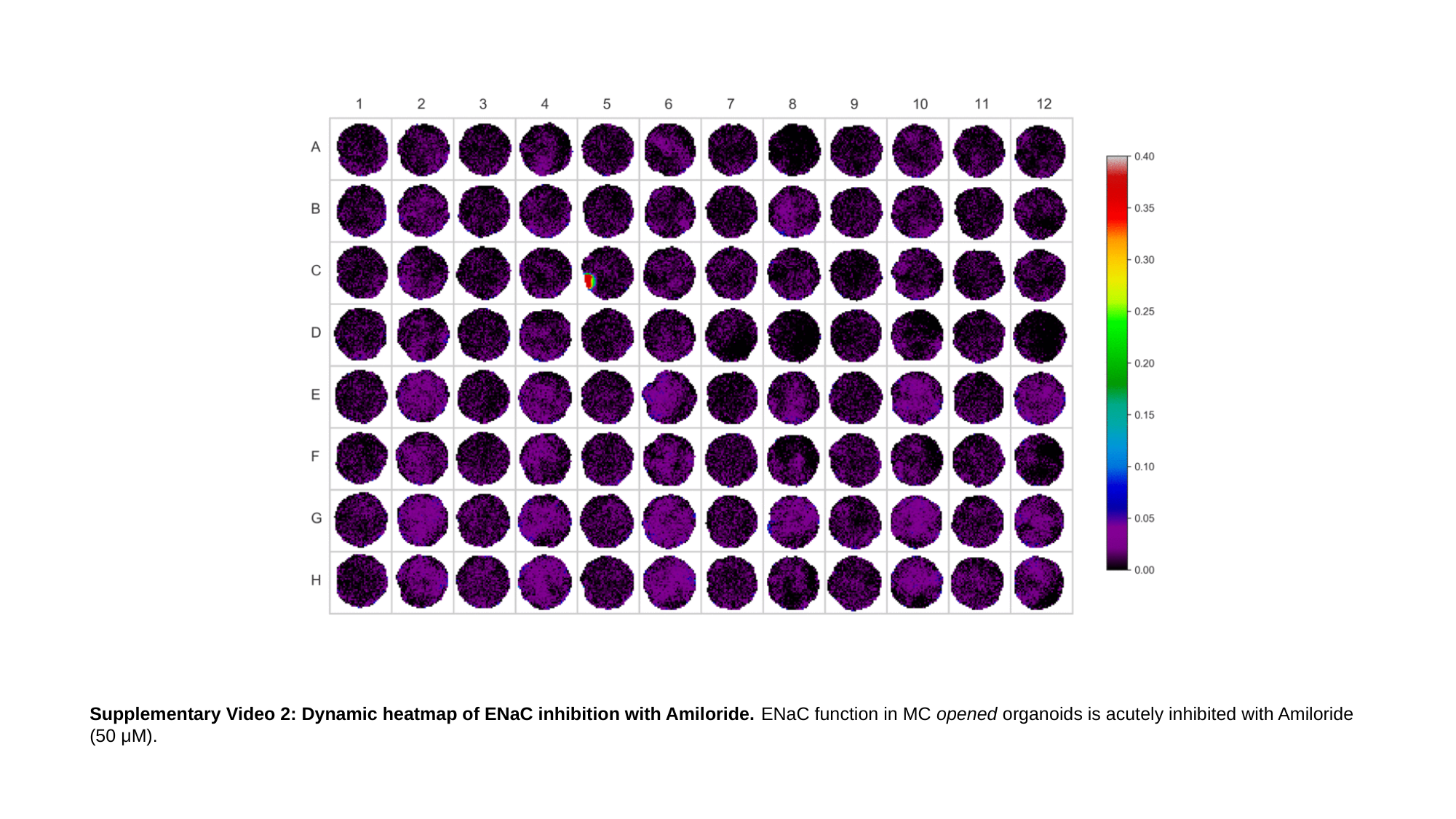

Supplementary Video 2: Dynamic heatmap of ENaC inhibition with Amiloride. ENaC function in MC opened organoids is acutely inhibited with Amiloride (50 μM).
